# The L-type voltage-gated Ca^2+^ channel CaV1.2 regulates fibroblast-like synoviocytes invasiveness during rheumatoid arthritis

**DOI:** 10.64898/2026.10.12.758622

**Authors:** Mark R. Tanner, R. Nicholas Elston, Madilyn J. Brandt, Brinkley A. Morse, Liang Sun, Shengwei Zou, Joseph M. Hyser, Frank T. Horrigan, Teresina Laragione, Pércio S. Gulko, Christine Beeton

## Abstract

Fibroblast-like synoviocytes (FLS) are major drivers of synovial tissue inflammation and damage during rheumatoid arthritis (RA). A hallmark of the aggressiveness of RA-FLS is their invasiveness but the molecular mechanisms that regulate this invasiveness remain poorly understood. The KCa1.1 (BK, Slo1, MaxiK, *KCNMA1*) potassium channel is a major regulator of RA-FLS pathogenicity and inhibiting KCa1.1 expression or its activity with selective blockers reduces RA-FLS invasiveness and reduces disease severity and joint damage in rat models of RA. KCa1.1 blockade induces a Ca^2+^ influx in RA-FLS whose intracellular signaling results in increased cell adhesion to the extracellular matrix and thus a reduction in cell migration. However, the identity of the Ca^2+^-permeable channel responsible for the Ca^2+^ influx has remained elusive. Here, we demonstrate the co-expression of KCa1.1 with the L-type voltage-gated CaV1.2 channels at the plasma membrane of primary human and rat FLS. Selective blocking of the CaV1.2 channel blunts the Ca^2+^ influx induced by KCa1.1 blockade and abrogates the effects of KCa1.1 blockade on RA-FLS invasiveness, migration, cell shape, and podoplanin expression. Whereas KCa1.1 blockers reduce disease severity in rat models of RA, treating rats with the collagen-induced arthritis model of RA with both blockers of KCa1.1 and CaV1.2 eliminates the therapeutic benefits of treatment with KCa1.1 blockers alone. Together, these data demonstrate a physical and functional coupling of KCa1.1 and CaV1.2 channels in FLS during RA and its rat model that regulate the invasive phenotype of the FLS through modulation of intracellular Ca^2+^ homeostasis.

**Significance Statement:** This study identifies the expression of CaV1.2 Ca^2+^ channels in primary fibroblast-like synoviocytes isolated from the joints of patients with rheumatoid arthritis. CaV1.2 channels are functionally coupled with KCa1.1 K^+^ channels to regulate the highly invasive phenotype of these synoviocytes, as blocking KCa1.1 induces a Ca^2+^ influx through opening of CaV1.2 that increases the adhesion of fibroblast-like synoviocytes and inhibits their migration. Conversely, blocking CaV1.2 negates the beneficial effects of KCa1.1 on the synoviocytes *in vitro* and abrogates the therapeutics effects of a KCa1.1 blocker in reducing clinical signs of collagen-induced arthritis.

## Introduction

Rheumatoid arthritis (RA) is a chronic inflammatory disease of the synovial joints. In addition to the pro-inflammatory immune response within the synovium characteristic of RA, resident joint cells, such as fibroblast-like synoviocytes (FLS), become activated during RA and contribute to tissue destruction. Under physiological conditions, FLS compose a layer of the synovial lining and secrete components of synovial fluid and maintain the extracellular matrix. However, during RA, FLS develop a highly invasive and destructive phenotype that results in formation of synovial hyperplasia and tissue damage (1). Although directly targeting FLS could serve as a mechanism to reduce disease severity without inducing general immunosuppression, which is a major side effect of currently used therapies, no therapeutics have yet been developed to specifically target aggressive FLS during RA.

We previously identified the KCa1.1 channel as the predominant potassium channel expressed by FLS and as a major regulator of FLS pathogenicity during RA and its animal models (2-7). Blocking KCa1.1 with selective modulators or reducing channel expression with siRNA reduces RA-FLS invasion, proliferation, and protease secretion *in vitro* (2, 4, 7). Furthermore, KCa1.1-selective blockers such as paxilline and iberiotoxin reduce disease severity and synovial tissue destruction in rat models of RA (3-5, 8). Blocking KCa1.1 also inhibits the migration of cancer cells *in vitro* (9). The mechanisms by which KCa1.1 block reduces RA-FLS and cancer cell migration are not fully elucidated. KCa1.1 blockade in RA-FLS induces a Ca^2+^ transient (7), leading to alterations in β1 integrin expression and activation, cell adhesion, and migration (10). However, the molecular identity of the channel responsible for the Ca^2+^ influx following KCa1.1 block has remained unknown. RA-FLS express multiple Ca^2+^-permeable transient receptor potential (TRP) channels that regulate intracellular Ca^2+^ homeostasis (11-13), but membrane depolarization induced by KCa1.1 block might not be sufficient to open a TRP channel in FLS. Indeed, some TRP channels expressed by FLS, including TRPC1 and TRPM7, are non-voltage gated. Some, such as TRPV1 or TRPV2, are weakly voltage-gated but the depolarization induced by KCa1.1 block would be insufficient to induce their opening in the absence of a primary activator such as temperature or capsaicin. Some TRP channels, such as TRPC5, are sensitive to depolarization but require a significant elevation of [Ca^2+^]i for opening. We therefore set out to identify the Ca^2+^-permeable ion channel responsible for the Ca^2+^ transient after KCa1.1 blockade.

In this study, we identified the expression of the L-type voltage-gated CaV1.2 Ca^2+^ channel at the plasma membrane of RA-FLS and demonstrated that inhibiting CaV1.2 prevents the KCa1.1 block-induced Ca^2+^ transient in RA-FLS. Furthermore, inhibition of CaV1.2 counters the effects of KCa1.1 block on RA-FLS invasiveness and negates the beneficial effect of inhibiting KCa1.1 in a rat model of RA. Taken together, these findings indicate a novel functional interaction between Cav1.2 and KCa1.1 in regulating RA-FLS.

## Results

### RA-FLS express functional CaV1.2 channels

We used a qRT-PCR screen to detect voltage-gated Ca^2+^ channel mRNA in RA-FLS and OA-FLS extracts and found that the transcripts of *CACNAC1*, the gene encoding the pore-forming α subunit of CaV1.2, were the most highly expressed in RA-FLS and OA-FLS (Fig. 1A). In addition, this expression was more than 2-fold higher in RA-FLS than OA-FLS.

**Figure 1.**
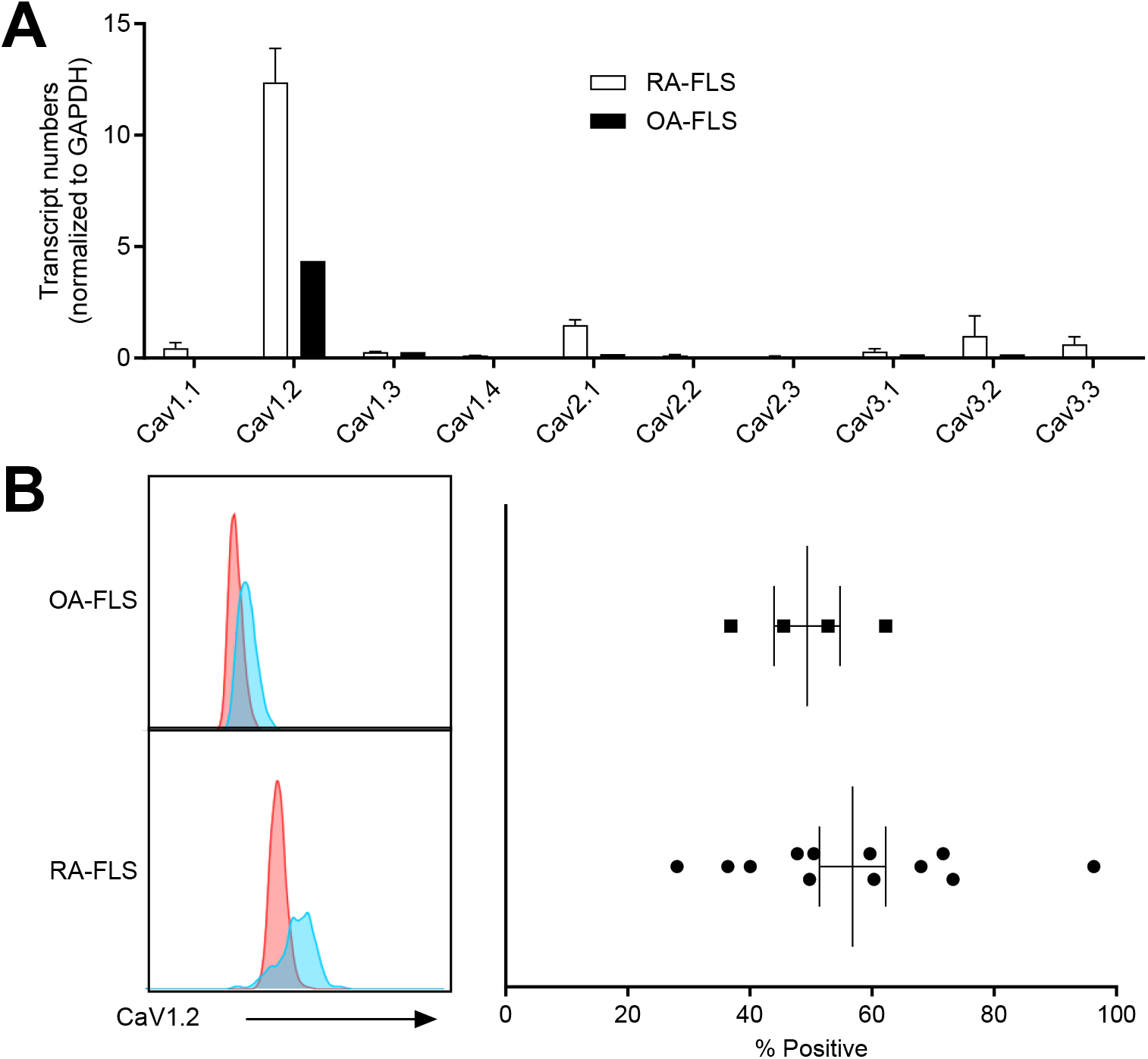
Primary human FLS express CaV1.2 channels at their plasma membrane. A, qRT-PCR-based quantification of transcripts of different voltage-gated calcium channels in RA- and OA-FLS. N=2 RA- and 1 OA-FLS donor. B, Left, example flow cytometry histograms of RA- and OA-FLS that were either unstained (red) or stained for the surface expression of CaV1.2 (blue). Right, quantifications of flow cytometry-based analyses of CaV1.2 surface expression in RA- and OA-FLS. N=12 RA-FLS and 4 OA-FLS.

Using flow cytometry and antibodies against the pore-forming α subunit of CaV1.2, we found that CaV1.2 is expressed at the protein level by both RA-FLS and OA-FLS, with 57 ± 5% of RA-FLS and 49 ± 5% of OA-FLS expressing the channel (Fig. 1B). HEK 293 cells expressing either CaV1.2 or CaV1.3 were used to validate the antibodies used in the flow cytometry assays (Supplemental Fig S1).

To determine whether CaV1.2 is expressed as a functional channel at the plasma membrane of RA-FLS, we used the whole-cell configuration of patch clamp electrophysiology and recorded inward Ca^2+^ currents. The inward current became detectable above -50 mV and reached a peak around 0 mV before decreasing progressively until 70 mV (Fig. 2A and B). Nifedipine is a well-characterized blocker of CaV1.2 (14) and the application of 1 µM nifedipine reduced current amplitudes recorded at 0 mV (Fig. 2C and D). Resting potential measurements from RA-FLS varied from -50 to -22 mV with a mean of -31.3 ± 1.9 mV (mean ± SD, N=6 cells) (Fig. 2E and F), near the threshold for CaV1.2 activation and similar to a previous measurement in rat FLS exposed to pro-inflammatory cytokines (-38.6 ± 2.8 mV) (15).

**Figure 2.**
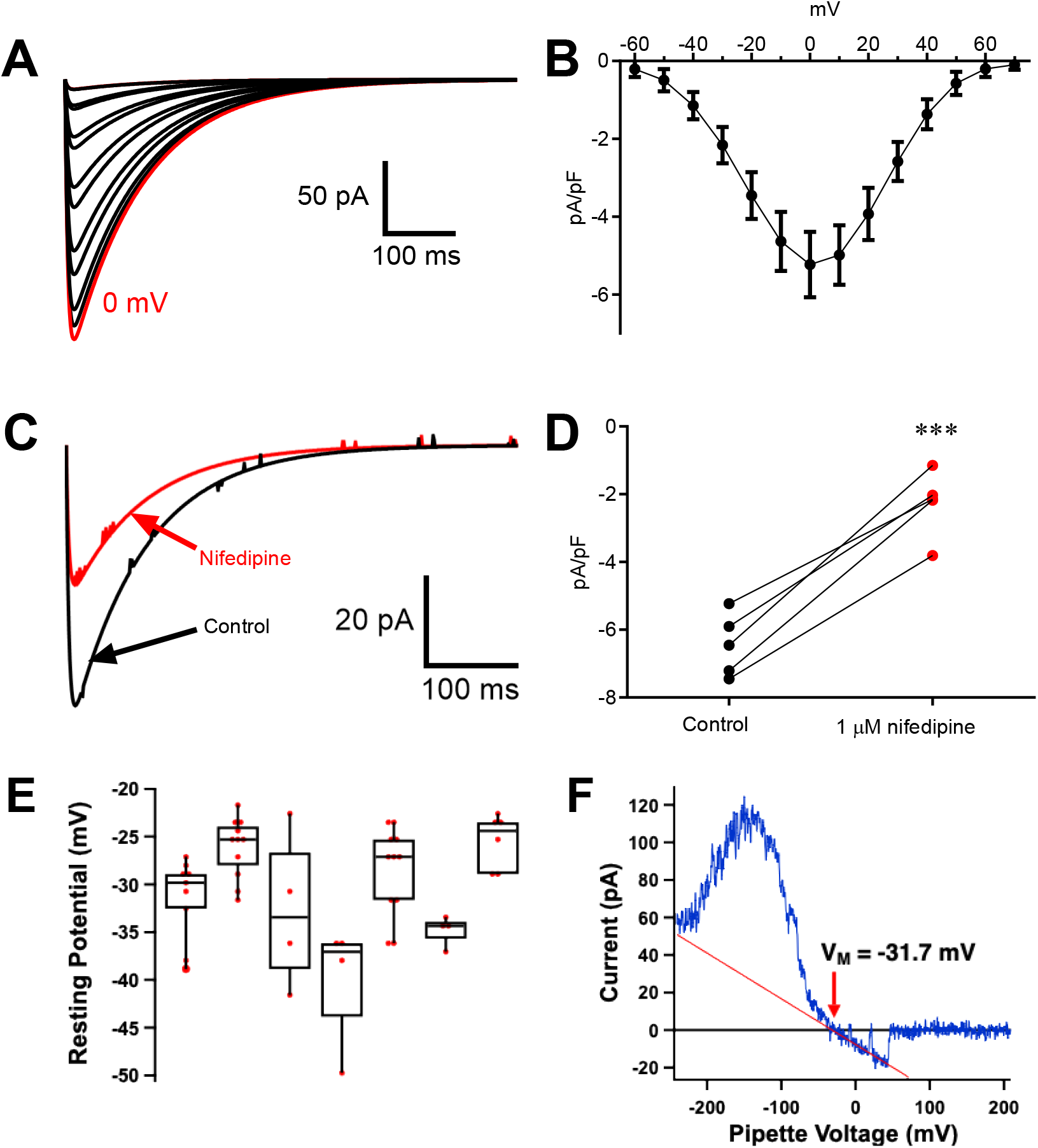
RA-FLS express functional CaV1.2 channels. A, Representative whole-cell recording of CaV current in an RA-FLS. B, The I-V plot of the data in C. C, Representative whole-cell recording of whole-cell CaV1.2 current in RA-FLS before (black) and after (red) addition of 1 µM nifedipine to the external solution. D, Quantification of the data in E. N=5. ***p<0.001 by paired t-test. E, Resting potential of 6 cells (N=4-10 measurements per cell). F, Attached patch recording of BK potassium current evoked by 10 ms voltage ramp from -250 to +200 mV (applied to pipette containing 140 mM K^+^) reverses at -31.7 mV corresponding to resting potential. Red line is fit to single channel current with slope conductance of 243 pS consistent with BK channel.

### CaV1.2 physically interact with KCa1.1 in RA-FLS

In vascular myocytes, KCa1.1 and Cav1.2 form a molecular complex with caveolin (16). We therefore investigated whether these two channels form a molecular complex in RA-FLS. Co-immunoprecipitation of the α subunits of CaV1.2 and KCa1.1 showed that the two channels physically interact in RA-FLS (Fig. 3).

**Figure 3.**
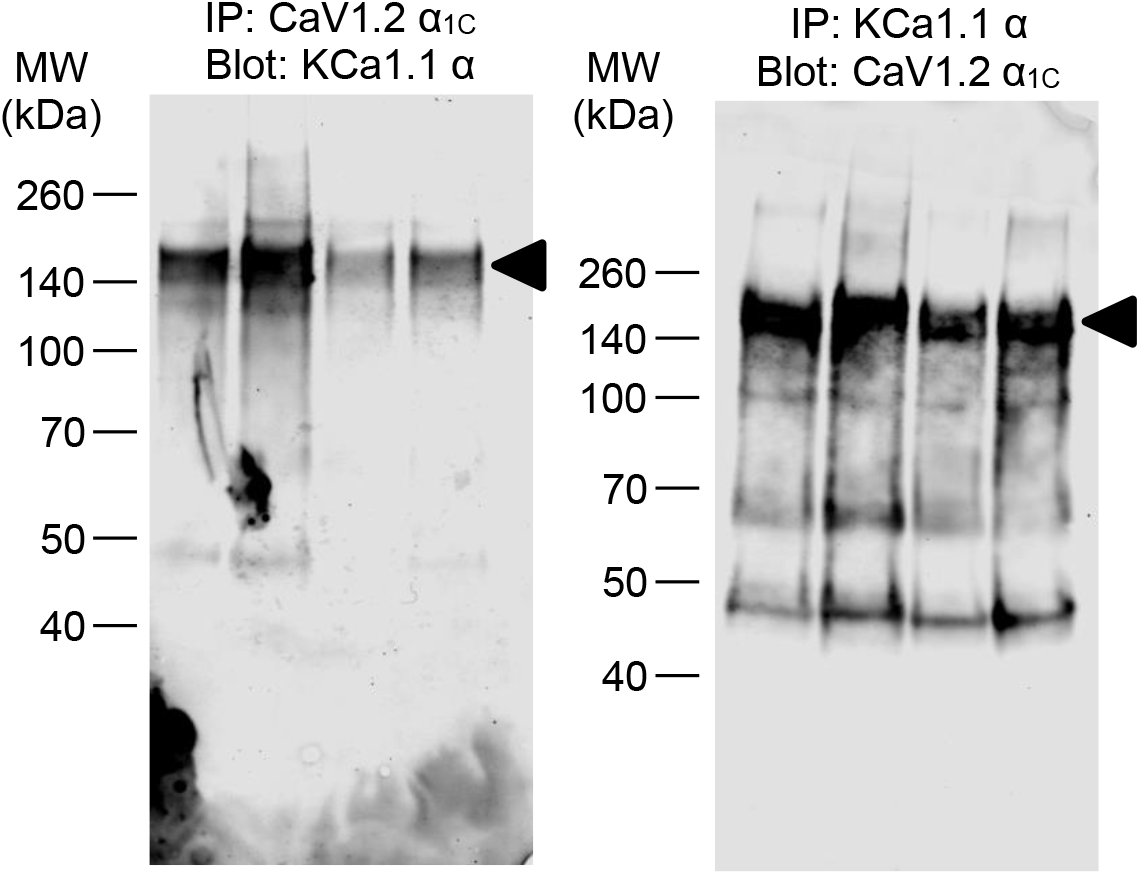
KCa1.1 and CaV1.2 physically interact in RA-FLS. Co-immunoprecipitation of CaV1.2 α1C and KCa1.1 α in protein extracts of RA-FLS from 4 different donors.

### CaV1.2 block reverses the KCa1.1 block-induced calcium transient in RA-FLS

Blocking KCa1.1 with paxilline induces a Ca^2+^ transient in RA-FLS (7). To determine if CaV1.2 is responsible for this Ca^2+^ transient, RA-FLS expressing the GCaMP6 calcium indicator, which increases in GFP fluorescence in the presence of intracellular calcium, and the fluorescent actin marker LifeAct-Ruby were treated with either paxilline alone to elicit the Ca^2+^ transient or with both paxilline and the CaV1.2 blocker nifedipine. As previously shown (7), blocking KCa1.1 with paxilline elicited a Ca^2+^ transient throughout the cytosol of most RA-FLS recorded, as shown through the large maximum GCaMP6 fluorescence increase following paxilline treatment (Fig. 4). In the presence of nifedipine, this paxilline-mediated Ca^2+^ transient was blunted, showing that the Ca^2+^ influx is mediated by CaV1.2.

**Figure 4.**
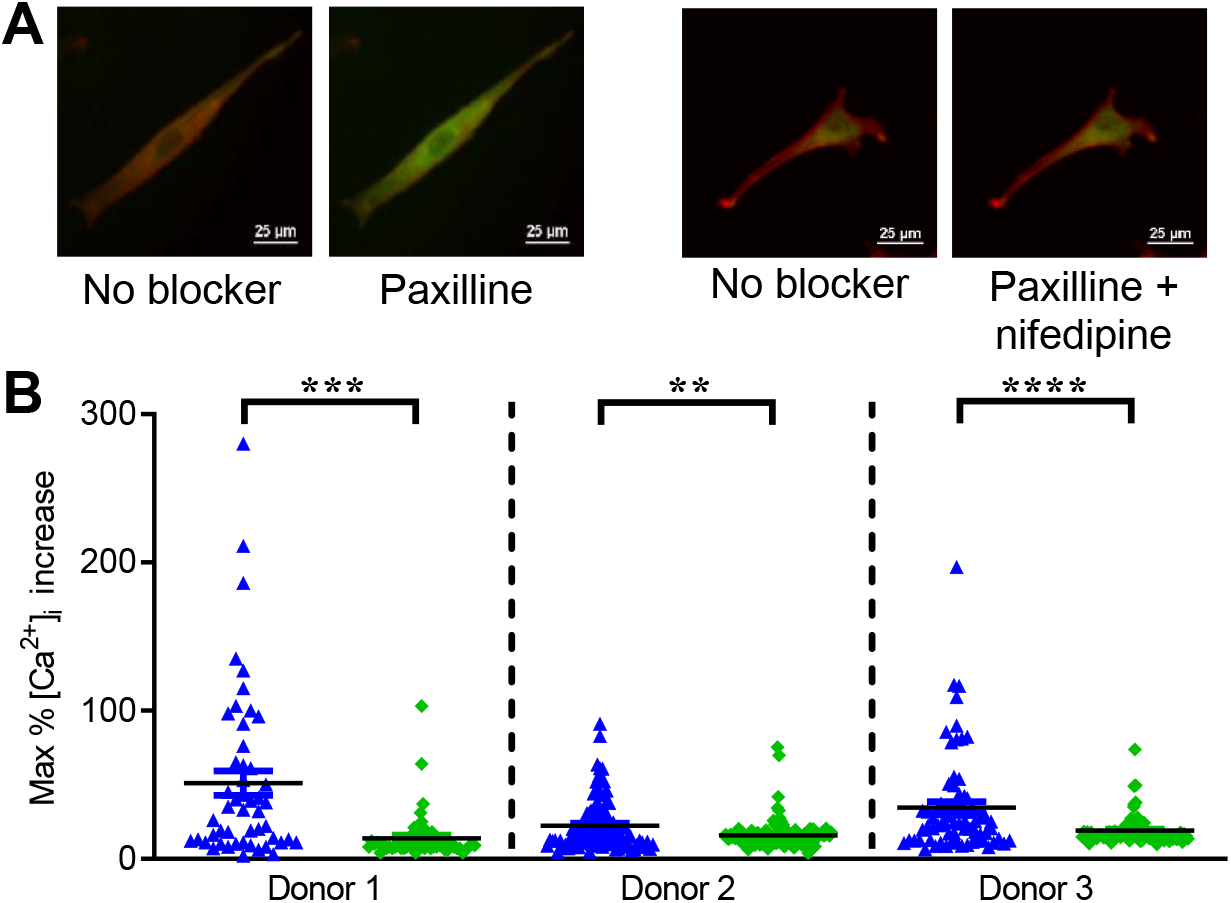
Blocking CaV1.2 blunts the paxilline-induced Ca^2+^ transient. A, representative images of RA-FLS expressing ruby-tagged actin (LifeAct-Ruby, red) and a cytosolic GFP Ca^2+^ sensor (GCaMP6, green in the presence of free cytosolic Ca^2+^) before blocker addition and after addition of paxilline alone or nifedipine + paxilline. Scale bars = 25 µm. B, quantification of the GCaMP6 Ca^2+^ sensor fluorescence 10 min after addition of paxilline alone 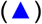 or both paxilline and nifedipine 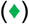. Each data point represents a different cell. **p<0.01, ***p<0.001, ****p<0.0001.

### CaV1.2 block reverses the effect of KCa1.1 blockers on RA-FLS invasiveness *in vitro*

KCa1.1 is a critical regulator of RA-FLS migration and invasion (2, 7, 10). Upon treatment with paxilline, RA-FLS invasiveness through Matrigel was reduced in half and this was reversed by the addition of nifedipine (Fig. 5A). Invasion through Matrigel requires the cells to migrate, alter their shape, and adhere to the extracellular matrix. To determine which of these aspects of invasiveness are regulated by KCa1.1 and CaV1.2, we assessed the migration of RA-FLS in an *in vitro* wound-healing assay. As previously shown (7), paxilline reduced the migration of RA-FLS, an effect not seen with nifedipine or the combination of both blockers (Fig. 5B,C). RA-FLS can be spindle-shaped or more rounded (Supplemental Fig. S2) and we quantified their circularity after treatment with the KCa1.1 and CaV1.2 blockers. After treatment with paxilline, RA-FLS exhibit an altered morphology, in which the percentage of cells in the classic fibroblastic spindle shape is reduced and have a higher circularity. However, treatment with nifedipine reduced this paxilline-induced change in cell shape (Fig. 5D). Podoplanin is a mucin-type transmembrane glycoprotein overexpressed by RA-FLS (17). It is coupled to the actin cytoskeleton and regulates cell shape and migration in a Ca^2+^-dependent manner (18, 19). We therefore assessed the effects of blocking KCa1.1 and CaV1.2 on its expression. Paxilline reduced the expression of podoplanin in almost half and this was normalized by the addition of nifedipine (Fig. 5E). Neither paxilline nor nifedipine had an effect on RA-FLS expression of cadherin-11 (Supplemental Fig. S3A), another adhesion protein known to regulate RA-FLS invasion (20), or on the expression of lubricin (Supplemental Fig. S3B), a component of joint lubrication secreted by FLS (21).

**Figure 5.**
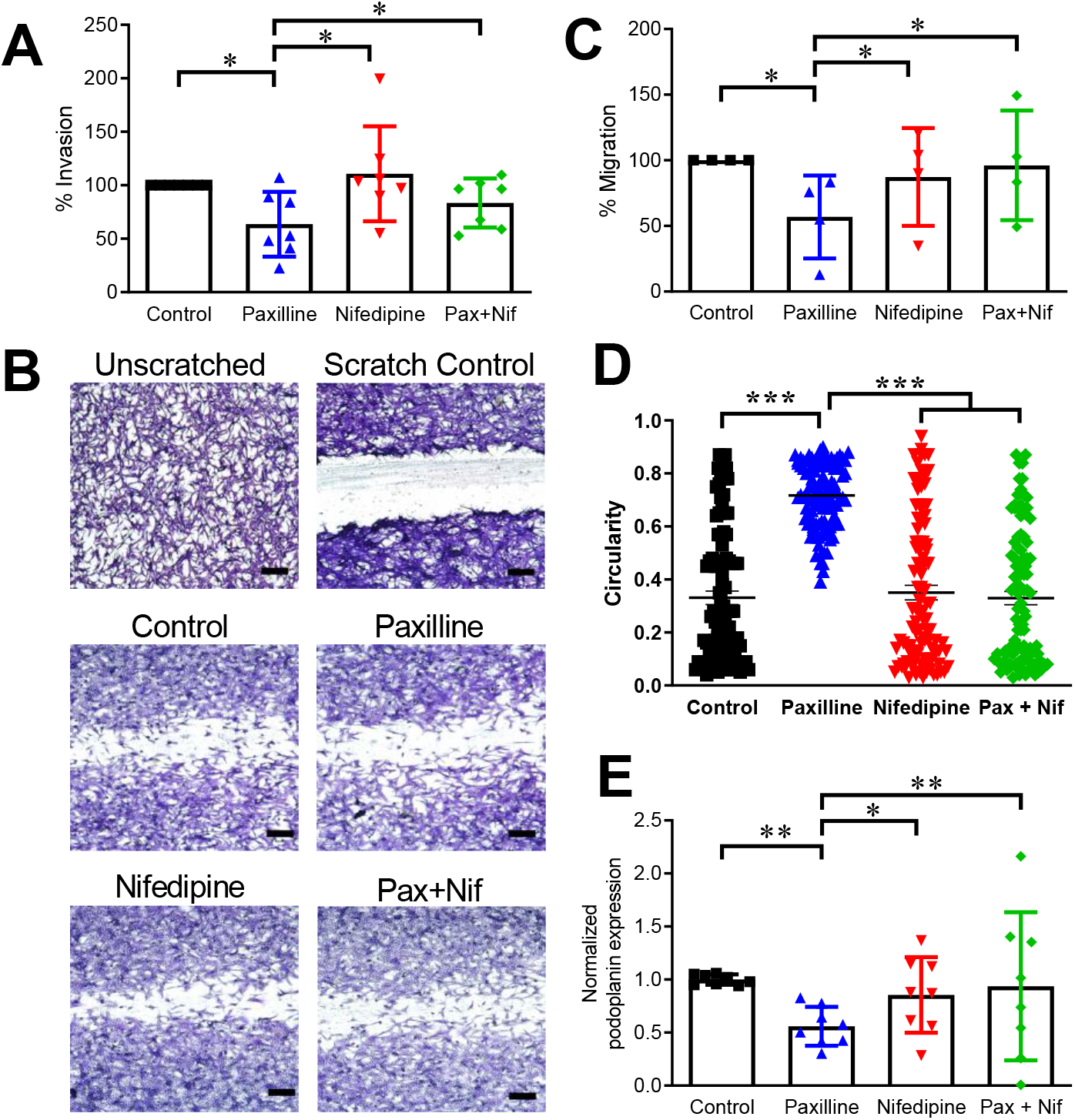
CaV1.2 block reverses the effect of KCa1.1 blockers on RA-FLS phenotypes *in vitro*. A, RA-FLS invasion through Matrigel-coated transwell inserts in the presence of media alone (■) 10 µM paxilline 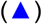, 1 µM nifedipine 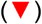, or both paxilline and nifedipine 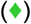. Each data point represents a different RA-FLS donor. B, representative images of RA-FLS migration into a scratched area in the presence or absence of 10 µM paxilline and 1 µM nifedipine. Scale bar=50 µm. C, Quantification of RA-FLS migration into a scratched area following treatment with 10 µM paxilline and 1 µM nifedipine. Each data point represents a different RA-FLS donor. D, Quantification of the circularity of RA-FLS after treatment with 10 µM paxilline and 1 µM nifedipine. N = 4 RA-FLS donors; each data point represents a different cell. E, Podoplanin expression normalized to that of control RA-FLS after treatment with 10 µM paxilline and 1 µM nifedipine. Each data point represents a different RA-FLS donor. *p<0.05, **p<0.01, ***p<0.001.

### CaV1.2 and KCa1.1 blockers alter disease severity in the collagen-induced arthritis (CIA) rat model of RA

We used the CIA rat model of RA to determine whether systemic treatment with nifedipine alters the effect of paxilline on disease severity. As previously shown (3-5), paxilline significantly reduced disease severity in this model. Surprisingly, we found that treatment with nifedipine alone significantly reduced disease severity in CIA (Fig. 6 and Supplemental Fig. S4). By contrast, the combination of paxilline and nifedipine eliminated the benefits of either compound alone as the course of disease in this group of rats was no different than that of rats treated with vehicle alone.

**Figure 6.**
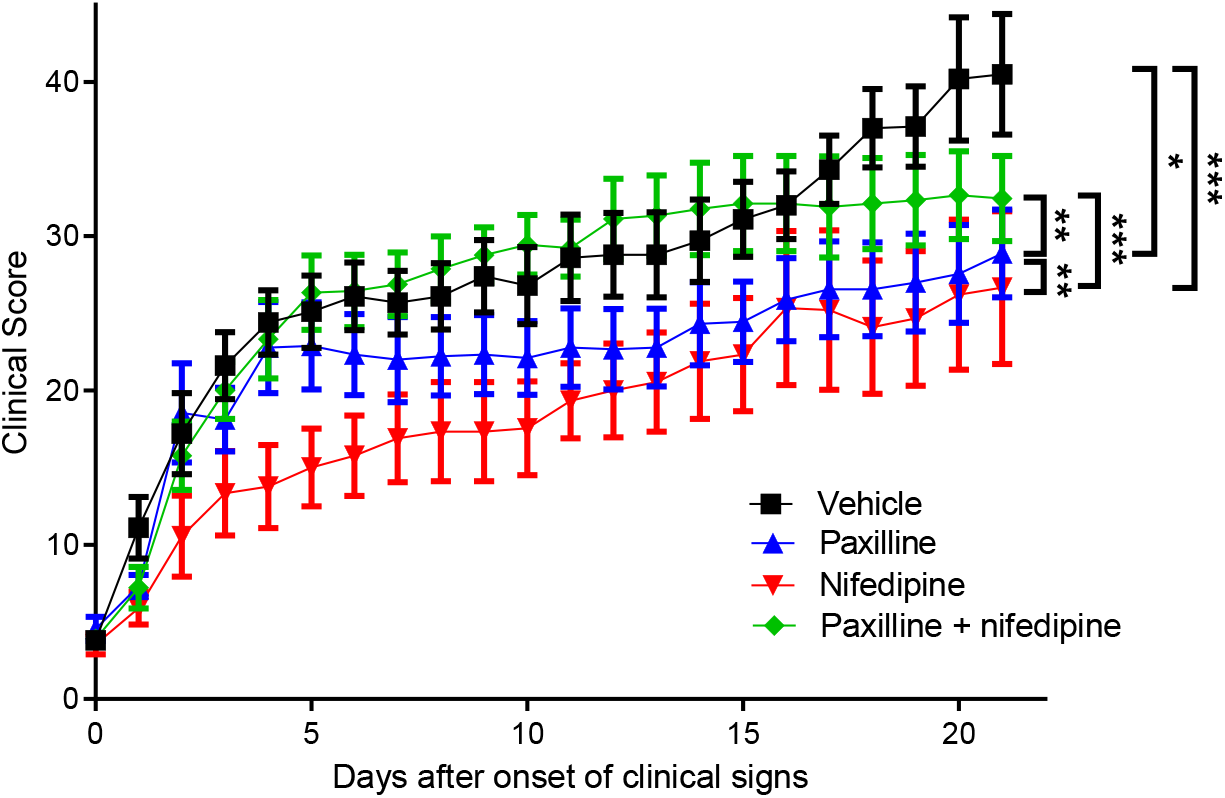
CaV1.2 and KCa1.1 blockers negate each other’s benefits in reducing disease severity in a rat model of RA. Clinical scores quantifying paw swelling of rats with CIA treated with vehicle (■), 20 mg/kg paxilline every-other-day 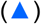, 0.1 mg/kg nifedipine every-other-day 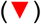, or both paxilline and nifedipine 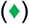. Mean ± SEM. N = 9-10 rats per group. Data from two independent assays were combined.

Following systemic treatment with nifedipine and paxilline, alone or in combination, for 21 days after disease onset, we isolated FLS from the paws of rats from each group and used flow cytometry to determine the percentage of cells expressing KCa1.1 and CaV1.2. Similar to human RA-FLS, CIA-FLS expressed both KCa1.1 and CaV1.2 and paxilline and nifedipine treatment did not alter expression of either channel (Supplemental Fig. S5).

We used joint histology and X-rays, and a standardized scoring system to quantify changes in disease severity at the tissue level between treatment groups. Rats with CIA and treated with vehicle had the most bone and cartilage damage, and synovial inflammation (Fig. 7A). Those rats treated with either paxilline or nifedipine had a marked reduction in cartilage erosions, synovial hyperplasia, pannus extensions, and immune infiltrates compared to vehicle-treated rats (Fig. 7B). Treatment with a combination of paxilline and nifedipine, however, had more bone and cartilage damage and synovial inflammation and were thus more similar to the vehicle-treated animals in those aspects of disease.

**Figure 7.**
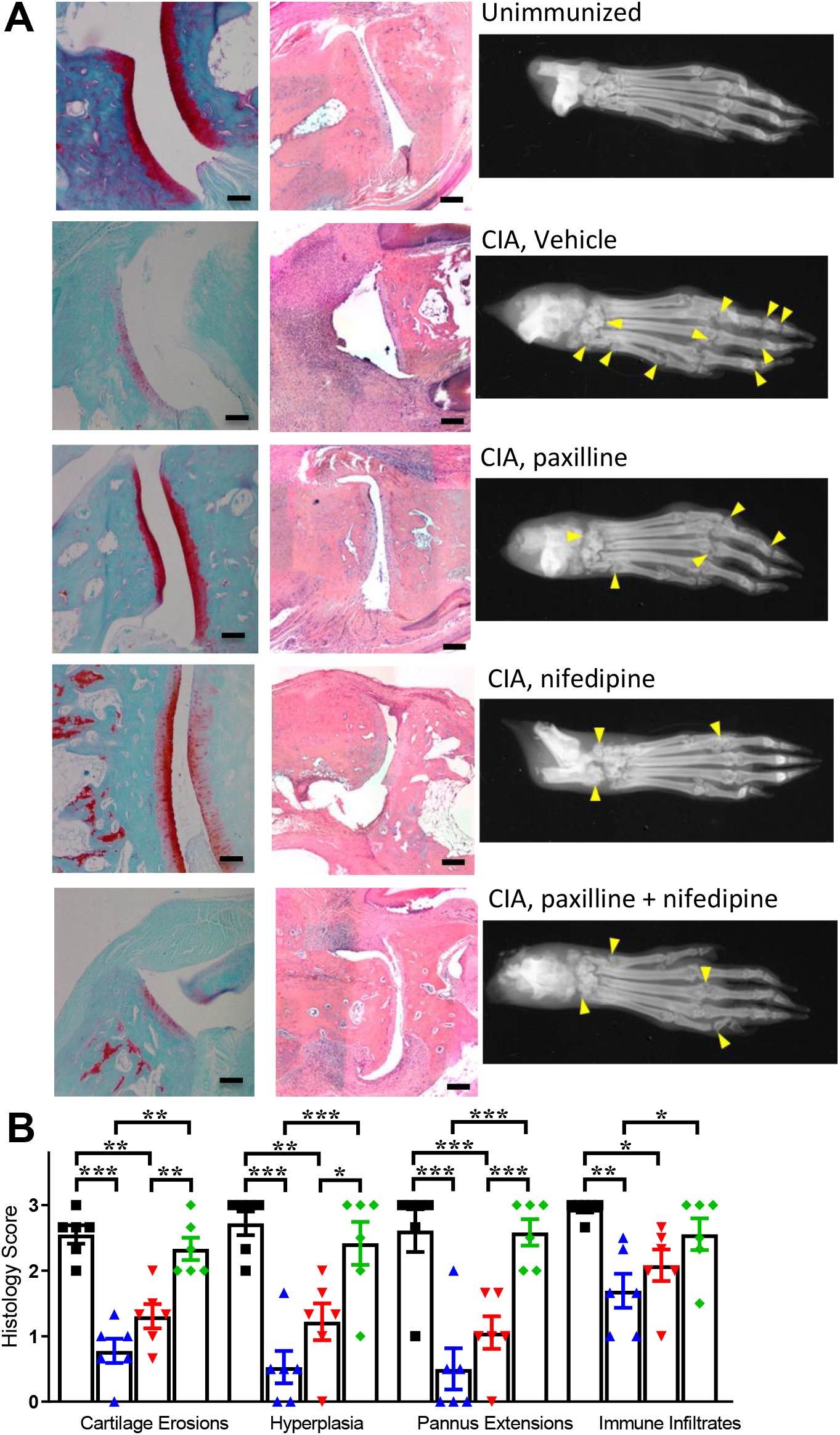
CaV1.2 and KCa1.1 blockers negate each other’s benefits in preventing joint damage in a rat model of RA. A, Left and middle, Representative images of joint sections from rats of each treatment group stained with Safranin O - fast green (left) or hematoxylin and eosin (middle). Scale bars = 100 µm. Right, representative X-rays of hind paws of rats from each treatment group. Yellow arrows indicate regions of noticeable bone damage. B, Quantification of histologic hallmarks of disease in CIA rats treated with vehicle (■), paxilline 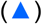, nifedipine 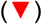, or both paxilline and nifedipine 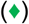. Each data point represents a different rat. *p<0.05, **p<0.01, ***p<0.001.

## Discussion

In this study, we identified an interplay between the KCa1.1 potassium channel and the CaV1.2 Ca^2+^ channel in RA-FLS. KCa1.1 has previously been proposed as a therapeutic target for RA as blocking this channel inhibits multiple pathogenic phenotypes of RA-FLS and KCa1.1 blockers reduce disease severity in multiple RA animal models (2-5, 7, 10). The mechanisms by which KCa1.1 regulates RA-FLS have remained largely unknown, though we previously found that KCa1.1 blockade induces a Ca^2+^ transient in RA-FLS that regulates β1 integrin expression and activation and thus cell adhesion to the extracellular matrix and migration. Here, we identified CaV1.2 as the predominant Ca^2+^ channel regulating this Ca^2+^ transient. This study is also the first to characterize functional CaV1.2 channels in RA-FLS.

This study is the first to show RA-FLS expression of functional CaV1.2 channels. While generally considered to be expressed by excitable cells, such as cardiomyocytes and neurons, CaV1.2 is also expressed by non-excitable cells, including other types of fibroblasts, such as those in human skin, rat tooth pulp, and gerbil inner ear (22-25). Further studies are needed to determine if CaV1.2 is expressed in other fibroblast types and what role it has in regulating fibroblast phenotypes in homeostatic conditions and during diseases with fibroblast involvement.

The more than two-fold higher expression of CaV1.2 α at the mRNA level was not recapitulated at the protein level as RA-FLS and OA-FLS expressed CaV1.2 α at similar rates. This discrepancy between mRNA and protein level expression is commonly seen in many cell types and can be explained by differences in synthesis and turnover rates (26). As the protein is the functional unit, our interpretation is that RA-FLS and OA-FLS do not differ in expression levels of the CaV1.2 channel.

The mechanism by which KCa1.1 block induces the opening of CaV1.2 has not been determined in FLS. One possibility is that the membrane depolarization induced by KCa1.1 block is sufficient to activated CaV1.2.

RA-FLS are not the first cell type in which CaV1.2 channels are found to regulate cell migration as blocking CaV1.2 has been shown to promote the migration of cancer cells (27, 28). Interestingly, as in RA-FLS, KCa1.1 is also a regulator of cancer cell migration (29) and RA-FLS are often described as having a tumor-like phenotype (30). Further work is needed to determine whether these two channels have the same functional interactions in regulating the migration and metastatic potential of cancer cells as we have described in RA-FLS.

FLS express multiple Ca^2+^-permeable channels, including TRP channels (11-13). Functional coupling of KCa1.1 and TRP channels have been reported in dorsal root ganglion neurons and in the paraventricular nucleus of the hypothalamus (31, 32). It is therefore possible that such interactions between KCa1.1 and TRP channels also occur in FLS. However, given that nifedipine, a CaV1.2 inhibitor (33), reduces KCa1.1-block-induced effects in RA-FLS, our results suggest that functional coupling between KCa1.1 and CaV1.2 is a major regulator of FLS phenotypes.

Our patch-clamp recordings show a rapid inactivation of the CaV1.2 current in RA-FLS. Whereas many studies of CaV1.2 currents are performed with Ba^2+^ as the charge carrier, we use solutions optimized for automated patch-clamp in which the charge carrier is Ca^2+^. Some splice variants of the α subunit of CaV1.2 exhibit Ca^2+^-dependent inactivation (34, 35), which is likely the reason of our observations. We have previously shown a switch in β regulatory subunits of KCa1.1 from β1 in OA-FLS to β3b in RA-FLS (2) and further studies are needed to determine whether OA-FLS and RA-FLS express the same or different splice variants of CaV1.2 α.

We have previously shown that, in addition to migration and invasiveness, KCa1.1 also regulates the proliferation and secretion of cytokines, proteases, and growth factors by RA-FLS (5, 7, 10). Here, we demonstrated that CaV1.2 mediates the beneficial effects of KCa1.1 blockade on RA-FLS migration and invasiveness and more work is necessary to determine the mechanisms by which KCa1.1 regulates RA-FLS proliferation and secretion and whether CaV1.2 is involved in regulating those phenotypes.

Nifedipine and other blockers of CaV1.2 and Cav1.3 are approved to treat hypertension and angina (36) and it would be interesting to determine whether patients with RA, treated with nifedipine or another CaV1.2 blocker, have a similar reduction in RA severity.

In assays with human FLS, nifedipine had no effect on RA-FLS invasiveness, migration, cell shape, or podoplanin expression, suggesting that the benefits of nifedipine in the rat model of CIA are not due to an effect on those aspects of RA-FLS mediated pathophysiology. However, blocking CaV1.2 could also affect other phenotypes of FLS, such as proliferation or protein secretion, to improve the clinical signs in CIA. In addition, CaV1.2 and Cav1.3 channels have been identified on multiple cell types besides FLS, including chondrocytes, osteoclasts, and macrophages and verapamil, another blocker of Cav1.2 and Cav1.3, treated a mouse model of RA, its beneficial effects attributed at least in part to action on macrophages (37-39). This benefit on macrophages is however overcome by the combined effects of paxilline and nifedipine on FLS. Further studies will be needed to fully define the CaV1.2 and Cav1.3 expression pattern and function in cells involved in the pathogenesis of RA during disease progression. As ion channel expression and function can also dramatically vary between species (40), a complete assessment across commonly used animal models of RA are also warranted.

Blocking both CaV1.2 and KCa1.1 negated the benefits from targeting either channel alone. Nifedipine and other selective blockers of CaV1.2 are used to treat hypertension and angina (41) and hypertension is frequently associated with RA (42). If blockers of KCa1.1 are ever tested in a clinical setting as potential therapeutics for RA or some cancers, it would be important to avoid the concomitant treatment of patients with a CaV1.2 blocker.

Overall, this study identifies a previously unknown expression and role for CaV1.2 in RA-FLS and demonstrates an interplay between CaV1.2 and KCa1.1 channels in regulating Ca^2+^ signaling, and thus the migration and invasiveness of these cells.

## Materials and Methods

### Origin and culture of cells

RA-FLS and OA-FLS were obtained as previously described (43, 44) under an IRB-approved protocol or purchased from Asterand. Diagnosis of RA or OA was defined in accordance with the criteria of the American College of Rheumatology (45). The characteristics of the subjects who donated FLS are in Supplemental Table 1. Cells were cultured in Dulbeco’s Modified Eagle Medium supplemented with 10% fetal bovine serum, 2 mg/ml glutamine, 2 mM sodium pyruvate, 10 units/ml penicillin, and 0.1 μg/ml streptomycin and used after the third passage.

HEK 293 cells expressing CaV1.2 or CaV1.3 (46) were a kind gift from Dr. Richard B. Silverman (Northwestern University, IL).

### Quantitative RT-PCR

RNA was isolated using the QIAGEN RNeasy Mini Kit. cDNA was generated using the iScript cDNA synthesis kit and 1 μg RNA and a 1:10 dilution of cDNA was used in each qPCR reaction. Primers used are listed in Supplemental Table 2. Target DNA was quantified using the Millipore SYBR Green PCR kit and BIO-RAD CFX96 realtime PCR system. Fold change and relative expression of all genes were calculated using the ΔΔCt method (47).

### Electrophysiology

Patch-clamp assays of Ca^2+^ current were conducted at room temperature (23-25ºC) in the whole-cell configuration using a Port-a-Patch system (Nanion Technologies, Livingston, NJ) connected to a HEKA EPC10-USB amplifier, and controlled with the PatchMaster software. The resistance of chips was 2-3.5 MΩ. The external solution contained 10 mM HEPES, 80 mM NaCl, 5 mM Glucose, 60 mM NMDG, 4 mM KCl 2mM CaCl2, 1 mM MgCl2, pH 7.4. The internal solution contained 10 mM EGTA, 10 mM HEPES, 110 mM CsF, 10 mM NaCl, 10 mM KCl, pH 7.2. CsF was used to block the overwhelming KCa1.1 currents in FLS (2, 4, 7). Cells were held at a resting potential of -80 mV and pulsed from -60 mV to 70 mV in 10 mV increments for 600 ms every 15 s. Nifedipine was diluted to 1 µM in external solution immediately before application to the cells.

The resting potential of RA-FLS was measured non-invasively in the attached patch configuration from the reversal potential of BK potassium current under the assumption, with 140 mM K^+^ in the pipette, that K^+^ current will reverse when the pipette voltage equals the resting potential (48). Currents were evoked by 10 ms voltage ramps (-250 to +200 mV pipette voltage) at room temperature (21–24°C) using an Axopatch 200B amplifier (Axon Instruments), PatchMaster acquisition software (HEKA), and Igor Pro (WaveMetrics, Inc., Lake Oswego, OR) for graphing and data analysis. Currents were filtered at 20 kHz with an 8-pole Bessel filter (Frequency Device, Inc.) and sampled at 200 kHz with an ITC-18 A/D converter (Instrutech). Pipette voltage was corrected for a +9.1 mV liquid junction potential, calculated according to the stationary Nernst–Planck equation (49) using LJPcalc. Pipettes were pulled from PG10150-4 glass (World Precision Instruments) and coated with wax (KERR Sticky Wax). The pipette solution contained (in mM) 140 K-methanesulfonic acid (K-MES), 2 MgCl2, 6 HCl, 20 HEPES pH 7.2 with MES. The bath solution contained 135 NaCl, 5 KCl, 2 CaCl2, 1 MgCl2, 10 HEPES, pH 7.2 w/NaOH.

### Flow cytometry

RA-FLS were treated for 72 hours with 10 μM paxilline and/or 1 μM nifedipine in 24-well plates. Cells were then detached from the plates with a cell lifter and stained with antibodies against cadherin-11 (Invitrogen, catalog 71-7600), total β1 integrin, which was conjugated to allophycocyanin (BD Pharmingen, catalog 559883), or activated β1 integrin, which was conjugated to phycoerythrin (BD Pharmingen, catalog 556049). For cadherin-11, cells were further stained with an anti-rabbit AlexaFluor-647 secondary antibody (Invitrogen, catalog A32733). FLS from healthy rats and those with CIA were fixed with formalin, permeabilized with 0.5% saponin, and stained with antibodies against KCa1.1α (Neuromab, catalog 75-022) or CaV1.2 α1C (Alomone labs, catalog ACC-003), followed by an anti-mouse IgG secondary antibody conjugated to AlexaFluor-647 (Invitrogen, catalog A32728). Data were acquired with a BD Canto II flow cytometer and analyzed using FlowJo software (Ashland, OR).

### Co-immunoprecipitations

The Capturem× IP & Co-IP Kit (Takara Bio) was used according to manufacturer instructions for RA-FLS lysis and immunoprecipitation of either KCa1.1α (Alomone catalog # APC-021) or CaV1.2 α1C (Alomone catalog # ACC-003). Samples were loaded onto NuPAGE 4-12% Bis-Tris gels and, after transfer, were probed for CaV1.2 α1C (Alomone catalog # ACC-003 at a 1:200 dilution) or KCa1.1α (Neuromab catalog # 75-022 at a 1:500 dilution). Secondary antibodies against rabbit IgG (Invitrogen catalog # A32735 at 0.05ug/mL) or mouse IgG (LiCOR catalog # 926-68070 at a 1:15,000 dilution) were used to detect the bands on a LiCOR Odyssey M.

### Calcium imaging

Lentivirus transduction was used to stably express LifeAct-Ruby and GCaMP6 in RA-FLS, as described previously (50, 51). Prior to imaging, cells were plated on 96-well flat-bottom plates overnight (1×10^3^ cells/well). Cell fluorescence was measured on a Nikon TiE inverted microscope before and after treatment with 1 μM nifedipine and/or 10 μM paxilline, with fluorescence measured every 2 seconds. Data was analyzed with Nikon Elements and Excel software and the peak intracellular calcium concentration was calculated as the maximum percentage GFP fluorescence change over baseline for each cell.

### Cell shape

RA-FLS were treated with 10 μM paxilline and/or 1 μM nifedipine and plated on 4-chamber glass slides. Cells were allowed to adhere for 16-18 hours before photos were taken using an AmScope MD35 camera and an Olympus BX41 microscope. The circularity of the cells was calculated with ImageJ (52), in which a value of 1 is a perfect circle and a value approaching 0 indicates an increasingly elongated polygon.

### Cell invasion

Cellular invasion was measured as described (2-5, 7), in which 50,000 RA-FLS were plated in the top compartment of Matrigel-coated Transwell inserts in the presence or absence of 10 μM paxilline and/or 1 μM nifedipine. Cells were allowed to invade for 16-18 hours and the cells that invaded through the Matrigel were fixed with formalin, stained with 0.5% crystal violet, and counted. Invasion of each condition was normalized to that of the number of invading cells in the control condition.

### Cell migration

Cellular migration was measured as described (7), in which 50,000 RA-FLS were plated in each chamber of 4-chamber slides and allowed to adhere. The cell monolayer was then scratched with a pipet tip and the cells were treated with 10 μM paxilline and/or 1 μM nifedipine. Migration into the scratched area was measured after 18 hours with data normalized to that of the migration of RA-FLS in the control condition. Images were taken using an AmScope MD35 camera and an Olympus BX41 microscope.

### Collagen-induced arthritis rat model

Female Lewis rats, age 8-11 weeks, were purchased from Envigo and housed in an AAALAC-accredited facility at Baylor College of Medicine under an IACUC-approved protocol. CIA was induced by a subcutaneous injection of 200 μl of a 1:1 emulsion of 2 mg/ml porcine type II collagen (Chondrex) and incomplete Freund’s adjuvant (Difco) (3-5). Seven days after the initial immunization, rats were given a booster of 100 μl of the emulsion. Disease onset was classified as when a rat developed at least one red or swollen toe or ankle joint. Clinical scores were determined on each rat daily as described (3-5, 53), in which a mildly swollen or red ankle was scored 2 points, a highly swollen or red ankle was scored 5 points, and each swollen or red toe joint was scored 1 point, for a maximum possible score of 60 per rat. Upon disease onset, rats were assigned to treatment groups, in which they received either 20 mg/kg paxilline intraperitoneally, 0.1 mg/kg nifedipine intraperitoneally, both paxilline and nifedipine, or vehicle (DMSO) every-other-day for twenty-one days. To avoid bias from differences in disease severity at start of treatment, we randomized rats to treatment groups by assigning every fourth rat that developed clinical signs of CIA on a given day to the same group (5). Following the completion of the study, rats were humanely euthanized prior to tissue collection.

### Histology and x-rays

Rat paws were fixed in formalin, decalcified, embedded in paraffin, and sectioned by the Human Tissue Acquisition and Pathology Core at Baylor College of Medicine, as described (3-5). Sections were stained for hematoxylin and eosin, imaged using Nikon Ci-L microscope in the Integrated Microscopy Core at Baylor College of Medicine, and scored as previously described (4). More details are provided in the supplement.

### Statistics

Comparisons between groups were completed with Mann-Whitney U tests. Clinical scores of CIA were compared using the nonparametric, repeated measures Friedmans tests with Dunn’s post-tests. All data are presented as mean ± standard error of the mean. Statistical analyses were completed using GraphPad Prism software.

## Acknowledgments

This work was supported by the Arthritis Foundation [Grant 6483] to CB and the National Institutes of Health (NIH) [Grant T32 HL007676] to MRT. The Cytometry and Cell Sorting, Mouse Phenotyping, Integrated Microscopy, and Human Tissue Acquisition and Pathology cores at Baylor College of Medicine are supported in part by funding from the NIH [Grants HG006348, RR007495, OD036336, DK56338, ES030285, and CA125123] and the Cancer Prevention and Research Institute of Texas [grants RP240432 and RP170719].

We thank Dr. Patrick Mumm (Nanion technologies, Munich) for providing us with the composition of the solutions optimized for the patch-clamp of CaV1.2 on automated systems.

