## Supplement for "The L-type voltage-gated Ca^2+^ channel CaV1.2 regulates fibroblast-like synoviocytes invasiveness during rheumatoid arthritis"

#### Supplementary information for:

### L-type voltage-gated $\text{Ca}^{2+}$ channels $\text{Ca}_v1.2$ regulate fibroblast-like synoviocytes invasiveness during rheumatoid arthritis

Authors: Mark R. Tanner<sup>1</sup>, R. Nicholas Elston<sup>1,2</sup>, Madilyn J. Brandt<sup>1,2</sup>, Brinkley A. Morse<sup>1,2</sup>, Liang Sun<sup>1</sup>, Shengwei Zou<sup>1</sup>, Joseph M. Hyser<sup>3</sup>, Frank T. Horrigan<sup>1</sup>, Teresina Laragione<sup>4</sup>, Pércio S. Gulko<sup>4</sup>, Christine Beeton<sup>1,5\*</sup>

<sup>1</sup> Department of Integrative Physiology, Baylor College of Medicine, Houston, Texas 77030, USA

<sup>2</sup> Graduate Program in Immunology and Microbiology, Baylor College of Medicine, Houston, Texas 77030, USA

<sup>3</sup> Department of Molecular Virology and Microbiology, Baylor College of Medicine, Houston, Texas 77030, USA

<sup>4</sup> Division of Rheumatology, Icahn School of Medicine at Mount Sinai, New York, New York 10029, USA

<sup>5</sup> Center for Translational Research on Inflammatory Diseases and Center for Drug Discovery, Baylor College of Medicine, Houston, Texas 77030, USA

\* Corresponding Author: Christine Beeton

#### Supplementary Information Text

##### Materials and Methods

###### Histology scoring

Sections of hind paws were stained with hematoxylin and eosin or with safranin O and fast Green, and scored by two investigators blinded to the animals' treatment group, using a well-established scoring system (1), as follows:

1. *Synovial inflammation*. Five fields were scored for the percentage of infiltrating mononuclear cells using the following scale: 0=absent; 1=mild (1–10%); 2=moderate (11–50%); 3=severe (51–100%). The mean of the five field for each animal was used for analyses.
2. *Synovial hyperplasia*. 0=absent; 1=mild (5–10 layers); 2=moderate (11–20 layers); 3=severe (>20 layers).
3. *Extension of pannus formation*. 0=absent; 1=mild; 2=moderate; 3=severe.
4. *Cartilage erosion*. Percentage of the cartilage surface that was eroded: 0=absent; 1=mild (1–10%); 2=moderate (11–50%); 3=severe (51–100%).

#### Supplementary Tables

| Donor | Diagnosis | Gender | Ethnicity | Age | Disease duration (years) | RF | Medications | Origin |
| --- | --- | --- | --- | --- | --- | --- | --- | --- |
| OA-1 | OA | Female | African American | 56 | 2 | NA | None | P.S.G.-knee |
| OA-2 | OA | Female | Caucasian | NA | NA | NA | NA | P.S.G.-hip |
| OA-3 | OA | Female | Caucasian | 65 | 5 | NA | None | P.S.G.-knee |
| OA-4 | OA | Male | Caucasian | 56 | 25 | NA | None | P.S.G.-knee |
| OA-5 | OA | Male | Caucasian | 52 | 78 | NA | None | P.S.G.-knee |
| RA-1 | RA | Female | African-American | 49 | 11 | + | Plaquenil, prednisone | P.S.G.-knee |
| RA-2 | RA | Female | Hispanic | 64 | >10 | + | Methotrexate, prednisone | P.S.G.-knee |
| RA-3 | RA | Female | Caucasian | 66 | 20 | + | Etanercept, Prednisone | P.S.G.-knee |
| RA-4 | RA | Female | Caucasian | 57 | 21 | + | Leflunomide, Etanercept, Prednisone | P.S.G.-knee |
| RA-5 | RA | Female | Caucasian | 71 | 3 | + | Prednisone, Methotrexate | P.S.G.-knee |
| RA-6 | RA | Female | Caucasian | 51 | 12 | + | NSAID, Prednisone | Asterand |
| RA-7 | RA | Female | Caucasian | 46 | 10 | - | Infliximab, Methotrexate, | P.S.G.-knee |
| RA-8 | RA | Female | Caucasian | 68 | 5 | + | Etanercept | P.S.G.-knee |
| RA-9 | RA | Male | Caucasian | 54 | >10 | + | Etanercept | P.S.G.-knee |
| RA-10 | RA | Male | Caucasian | 79 | 15 | + | NSAID, Prednisone, Adalimumab | P.S.G.-hip |
| RA-11 | RA | Male | Hispanic | 48 | <1 | + | Prednisone, DMARD | Asterand |

**Table S1. Characteristics of the subjects who donated FLS for this study.** DMARD: disease-modifying anti-rheumatic drug; NSAID: non-steroidal anti-inflammatory drug; P.S.G., Dr. P.S. Gulko's group; RF: rheumatoid factor.

| Protein | Gene | NCBI gene number | GenBank accession number | Primer sequence |
| --- | --- | --- | --- | --- |
| Cav1.1 | CACNA1S | 779 | NM_000069 | GACCACTCAGAGCCATCAACAG<br>GGTAGTGACCAGCACGATGTTC |
| Cav1.2 | CACNA1C | 775 | NM_000719 | GCAGGAGTACAAGAACTGTGAGC<br>CGAAGTAGGTGGAGTTGACCAC |
| Cav1.3 | CACNA1D | 776 | NM_000720 | CTTCGACAACGTCCTCTCTGCT<br>GCCGATGTTCTCTCCATTCGAG |
| Cav1.4 | CACNA1F | 778 | NM_005183 | GAGGAAGTCCCTGATCGGCTTTC<br>CCCTCTGAGATAAGCACAGCCTC |
| Cav2.1 | CACNA1A | 773 | NM_000068 | CTGGTAGCCTTTGCCTTCACTG<br>ACACAGCCTTGAGCTTTGGCAG |
| Cav2.2 | CACNA1B | 774 | NM_000718 | TGGACCTGGAAAGCCAAGCAGA<br>GGTCACGATGTAGTGGCAGAAG |
| Cav2.3 | CACNA1E | 777 | NM_000721 | AGCGTGAGACAGGCAAAGCCAT<br>GGATGCACATCTCAAAGTAGCGC |
| Cav3.1 | CACNA1G | 8913 | NM_018896 | TTCACCGCAGTCTTTCTGGCTG<br>TGACGGAGATGAGCACCAACAG |
| Cav3.2 | CACNA1H | 8912 | NM_021098 | GGAACATCTCCACCAAGGCACA<br>TCCATCCTTGGATGACAGCACG |
| Cav3.3 | CACNA1I | 8911 | NM_021096 | GTGTCCAACTACATCTTCACGGC<br>GACAAGAAAGCCATCCAGCACG |

**Table S2. Primers used for qRT-PCR.**

#### Supplementary Figures

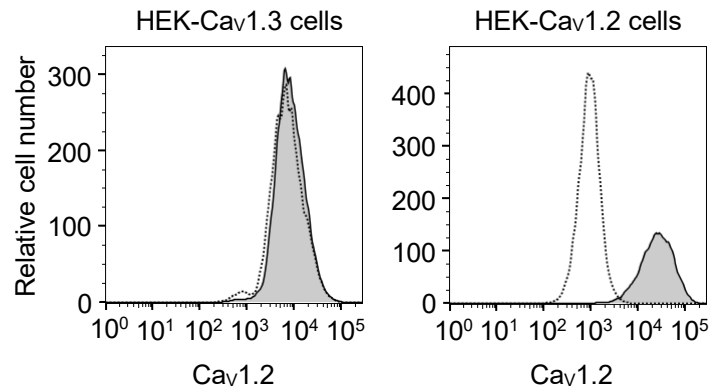

**Figure S1. Authentication of the anti-Cav1.2 $\alpha$ antibodies.** Flow cytometric histograms of HEK 293 cells stably expressing CaV1.3 (left) or CaV1.2 (right) stained for the expression of Cav1.2 $\alpha_{1C}$. Unstained cells are shown as dashed lines with clear histograms and the stained cells are shown as solid lines filled in gray.

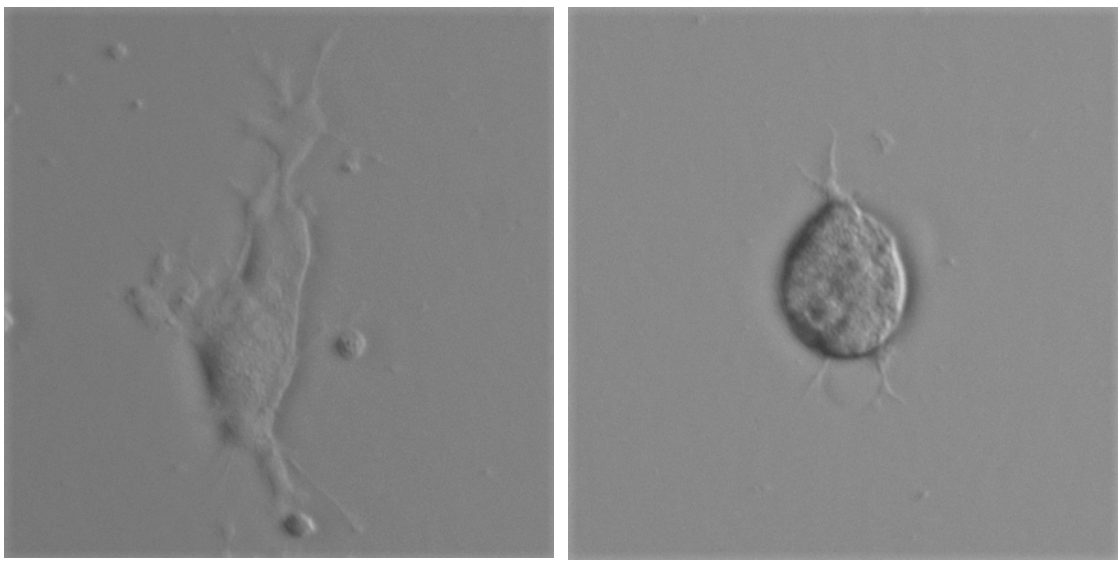

**Figure S2. RA-FLS shapes.** Representative images of RA-FLS with an elongated and irregular cell shape (left; circularity 0.07) and with a rounded shape (right, circularity 0.89).

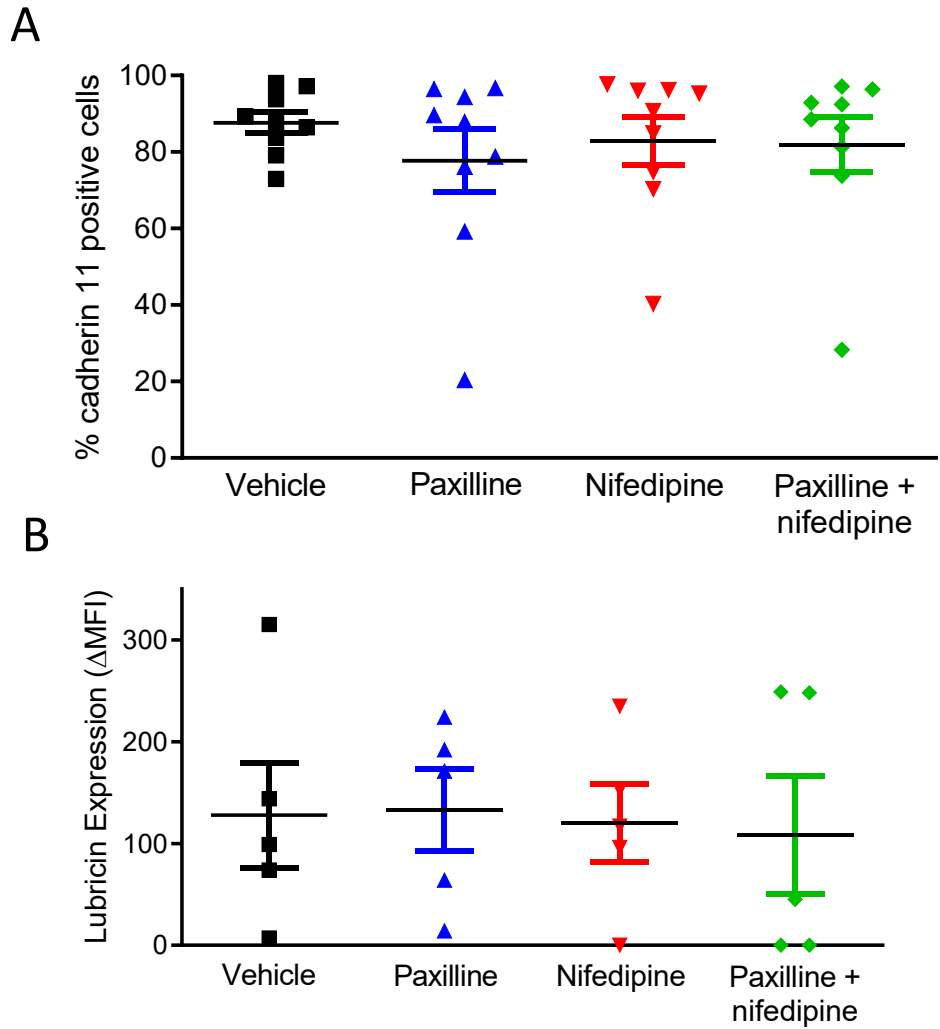

**Figure S3. No effect of KCa1.1 and CaV1.2 block on RA-FLS cadherin-11 and lubricin expression.**

A, RA-FLS surface expression of cadherin-11 following treatment with 10 μM paxilline and/or 1 μM nifedipine. B, RA-FLS expression of lubricin following treatment with 10 μM paxilline and/or 1 μM nifedipine. Each data point represents a different RA-FLS donor.

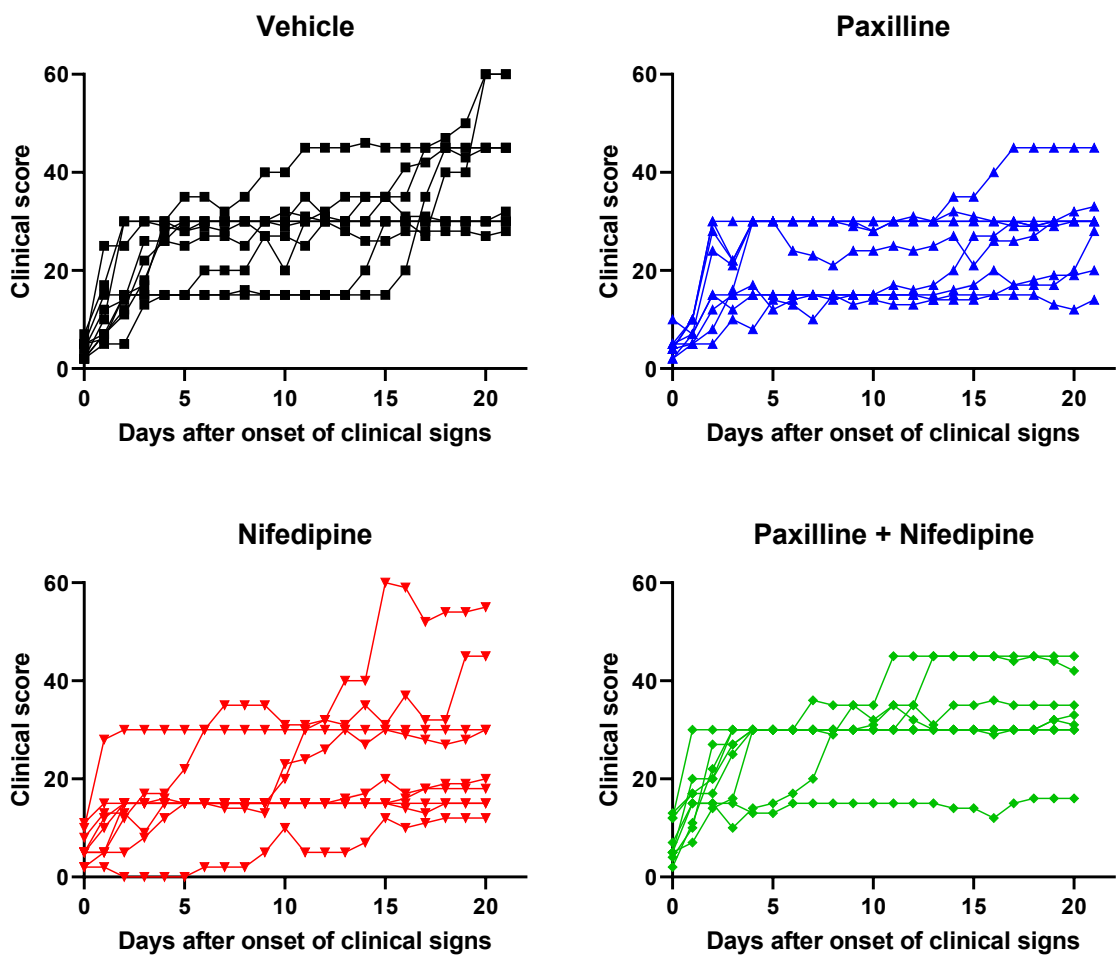

**Figure S4. Individual clinical scores of rats in the CIA trials.** Clinical scores quantifying paw swelling of rats with CIA treated with vehicle (■), 20 mg/kg paxilline every-other-day (▲), 0.1 mg/kg nifedipine every-other-day (▼), or both paxilline and nifedipine (◆). Each curve represents a single rat. Data from two independent assays were combined.

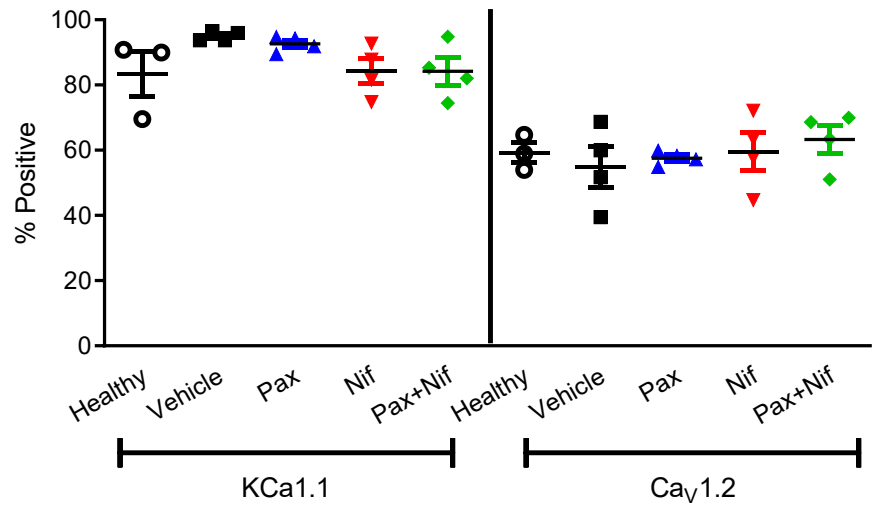

**Figure S5.** KCa1.1 and CaV1.2 expression in FLS isolated from healthy rats and rats with CIA treated with vehicle, paxilline, nifedipine, or both paxilline and nifedipine for 21 days prior to cell isolation. Each data point represents a different rat.

#### Supplementary Information References

1. M. Brenner *et al.*, Identification of two new arthritis severity loci that regulate levels of autoantibodies, interleukin-1β, and joint damage in pristane- and collagen-induced arthritis. *Arthritis Rheum* **64**, 1369-1378 (2012).
